# Contextual Biomedical Language Models for Imbalance-Aware Drug–Food Interaction Classification

**DOI:** 10.64898/2026.10.19.760076

**Authors:** Mahendar Yadav, Nalgonda Lokesh, Mohammed Zubair, Mansoor

## Abstract

Drug–food interactions (DFIs) are essential to patient safety because they can alter drug effectiveness or cause side effects. Automated classification of DFIs is difficult because of unstructured textual data, semantic ambiguity, and class imbalance, especially when separating neutral interactions. This paper presents an imbalance-aware deep learning approach for classifying drug–food interactions into three classes: safe, neutral, and unsafe. A structured dataset is created from textual drug interaction descriptions by extracting drug–food pairs and labeling them using rule-based heuristics. To address class imbalance, we apply class weighting and focal loss while fine-tuning the context-aware BioBERT language model. Our experiments show that the approach reaches an accuracy of 85% and a macro-averaged F1-score of 0.77, with a notable improvement on the minority neutral class (F1-score of 0.56). The main novelty of this work is the combination of contextual biomedical language modeling with imbalance-aware deep learning, enabling more effective classification of DFI datasets.

## 1 Introduction

Drug–food interactions (DFIs) are an important yet understudied factor affecting patient health. Food consumption can alter drug pharmacokinetics and pharmacodynamics, leading to reduced efficacy, undesirable side effects, or even toxic reactions. Grapefruit juice, for example, interferes with drug metabolism, and high amounts of vitamin K can neutralize the effect of anticoagulants. With the growing number of complex drug regimens and over-the-counter dietary supplements, DFIs have become an integral part of clinical decision-making.

Owing to the absence of automated solutions, identifying and classifying DFIs in clinical practice remains an open challenge. Information on DFIs mostly appears in unstructured form in drug labels, medical literature, and clinical guidelines. Manual extraction and interpretation of this knowledge is time-consuming and impractical in real-world settings. There is therefore a need for automatic classification of DFIs as safe, neutral, or unsafe with respect to drug effect.

Transformer architectures such as BERT and its biomedical counterpart BioBERT have proved effective in capturing contextual information in biomedical text. Such models provide deep contextual embeddings that support accurate analysis of domain-specific text. However, applying them directly to DFI classification raises several challenges. One is the lack of datasets for DFI classification; raw interaction descriptions often need to be converted into a structured representation using heuristic rules.

Another challenge is class imbalance. In realistic scenarios, the underrepresented neutral class is overshadowed by safe or unsafe interactions during training, causing biased learning. Accurate prediction of neutral interactions is further complicated by their ambiguity, as they overlap with characteristics of both safe and unsafe interactions. Machine learning models, including deep architectures, often perform poorly under such imbalance.

Existing approaches to interaction classification are mostly based on classical machine learning algorithms such as SVM and Random Forests, using handcrafted features such as TF-IDF vectors. These outperform many purely statistical models but cannot capture deeper information within biomedical text. Recent deep learning approaches address representation learning but remain ineffective at handling class imbalance and minority-class prediction. Moreover, most related studies address drug–drug interactions rather than drug–food interactions.

This paper proposes an imbalance-aware framework for classifying drug–food interactions with contextual biomedical language models. A structured dataset is generated by extracting drug–food interactions from unstructured text and applying rule-based labeling. The problem is formulated as a sequence classification task, and BioBERT is fine-tuned for it. An imbalance-aware objective function is used to improve minority-class performance, and leakage-free drug-level evaluation is employed to assess the results.

The complete source code and implementation details are available at https://github.com/NalgondaLokesh/Drug-Food-Interaction-BioBert.

To the best of our knowledge, this is the first work that applies BioBERT with imbalance-aware training to drug–food interaction classification.

The main contributions of this paper are:

1. A novel solution for DFI classification using contextual language models.
2. An automatic dataset construction pipeline that produces structured drug–food interaction pairs with corresponding labels.
3. An imbalance-aware training strategy combining focal loss and class weighting to improve minority-class prediction.
4. A leakage-free, drug-level evaluation strategy.
5. Experiments showing improved classification performance, particularly for the neutral class.

## 2 Related Work

Previous research on DFIs has primarily focused on elucidating their pharmacological and clinical nature and their effect on drug efficacy and patient health. For example, Bushra et al. conducted an extensive survey of food–drug interactions, discussing the effect of nutrition on the bioavailability of pharmaceutical compounds [1]. Despite its valuable insights, this work is descriptive, and computational techniques are needed to predict DFIs at scale. With advances in computing, machine learning became a focus for drug interaction prediction. Ryu et al. developed DeepDDI, a framework based on neural networks and structural similarity profiles of drug molecules [2], which predicts drug interactions and generates human-readable descriptions of interaction types. However, DeepDDI relies on structural features of molecules and does not specifically address drug–food interactions, since its training data consist of drug–drug interaction cases.

More recently, Kha et al. proposed an explainable machine learning approach for DFI prediction based on features extracted from chemical structures in the DrugBank and FooDB databases [3]. The approach used several classifiers, including XGBoost, and a feature selection technique that further improved accuracy. Although it performs well and explains its decisions, it depends on manually created chemical descriptors that cannot reflect the contextual relations contained in unstructured text.

Traditionally, Support Vector Machines (SVM) and Random Forests (RF) have been used for drug interaction prediction. These algorithms depend on handcrafted features such as molecular descriptors or TF-IDF text representations. Although they are reliable baselines, they struggle to incorporate the complex contextual and semantic relations present in biomedical text.

Recently, transformer-based models have significantly advanced natural language processing. Devlin et al. introduced BERT, a bidirectional transformer that learns deep contextual representations by jointly conditioning on left and right context using masked language modeling and next sentence prediction [4]. BERT can be fine-tuned for a wide range of downstream tasks with minimal architectural modification. Building on this, Lee et al. proposed BioBERT, a domain-specific extension of BERT pre-trained on large-scale biomedical corpora such as PubMed and PMC articles [5]. BioBERT substantially improves performance on biomedical text mining tasks, including named entity recognition, relation extraction, and question answering.

Despite the success of transformer-based models, their application to drug–food interaction prediction remains limited, particularly on imbalanced datasets. Class imbalance is a critical issue in DFI prediction because some classes, in particular the neutral class, are far less represented than others. Although this problem has been addressed in the general machine learning literature through class weighting and focal loss, these methods have not been discussed in DFI prediction studies.

Thus, prior work has produced models for DFI prediction, but with several limitations: most use structured chemical features and ignore textual information, DFI classification is less researched than DDI classification, and few works address class imbalance in biomedical data.

## 3 Methodology

This section describes the proposed methodology for classifying drug–food interactions with contextual biomedical language models. The approach comprises dataset preparation from unstructured sources, preprocessing, transformer-based modeling, handling of class imbalance, and comparison with classical machine learning methods.

### 3.1 Dataset Construction

A major limitation in DFI studies is the lack of structured, labeled datasets. To address this, a dataset was created from raw descriptions of drug–food interactions. The dataset was generated from a CSV file containing drug names and their food interaction descriptions [6].

Table 1 presents the dataset statistics. The raw dataset contains 1,988 drug–food interaction pairs after initial processing. The class distribution is strongly imbalanced: the neutral class comprises only 133 samples (6.7%), compared with 926 safe samples (46.6%) and 929 unsafe samples (46.7%). To address this imbalance, random oversampling was applied to create a balanced training set of 2,787 samples with 929 samples per class. The test set consists of 502 samples and retains the original class distribution for fair evaluation.

**Table 1.** Dataset statistics before and after resampling.

| Class | Training (before) | Training (after) | Test |
| --- | --- | --- | --- |
| Safe (0) | 926 (46.6%) | 929 (33.3%) | 242 (48.2%) |
| Neutral (1) | 133 (6.7%) | 929 (33.3%) | 38 (7.6%) |
| Unsafe (2) | 929 (46.7%) | 929 (33.3%) | 222 (44.2%) |
| **Total** | **1,988** | **2,787** | **502** |

Dataset preparation begins by extracting drug–food information from textual descriptions. First, the text was searched for keywords related to known food interaction terms (e.g., alcohol, grapefruit, milk, caffeine); when found, the food entities were labeled. Linguistic NER tools were then applied to extract domain-specific food items.

After extraction, the pairs were labeled using a rule-based approach:

- Texts with “Avoid,” “Contraindicated,” or “Danger” warnings are classified as **Unsafe**.
- Texts with “Limit,” “Monitor,” or “Use carefully” indications are classified as **Neutral**.
- All other texts are classified as **Safe**.

The resulting dataset consists of triplets of the form (*drug, food, label*). Duplicates were removed. The class distribution is not uniform, as there are fewer neutral interaction pairs than safe and unsafe pairs.

### 3.2 Data Preprocessing

Textual data was cleaned by removing special characters, extra whitespace, and other noise, and all text was converted to lowercase. Structured inputs were created using the template:

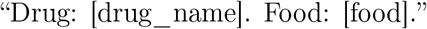

This input structure provides semantic context for classification.

To ensure proper evaluation and avoid leakage, drug-level splitting was used instead of conventional sample-based splitting. Drug entities are not split randomly across the train, validation, and test sets; all samples of a given drug belong to exactly one of them.

For transformer-based models, texts were tokenized with the BioBERT WordPiece tokenizer, which handles biomedical terms. All sequences were padded or truncated to a fixed maximum length.

### 3.3 Model Architecture

#### 3.3.1 BioBERT Model

The proposed architecture uses BioBERT, a Transformer model pre-trained on biomedical corpora, which generates contextual embeddings and learns semantic relationships between texts. A special classification token ([CLS]) is prepended to every input, and the final hidden-layer embedding of this token serves as a summary of the whole sequence. A classifier layer is added on top, and its outputs are passed through a softmax function to produce probabilities over three classes (safe, neutral, unsafe).

#### 3.3.2 Baseline Models

Two classical machine learning baselines were implemented:

- **TF-IDF + Support Vector Machine (SVM):** text is transformed into TF-IDF vectors and classified with an SVM.
- **TF-IDF + Random Forest:** the same features are classified with a Random Forest.

### 3.4 Handling Class Imbalance

The neutral class is strongly under-represented, biasing the dataset toward safe and unsafe classes. To address this, an imbalance-aware training technique based on focal loss was used:

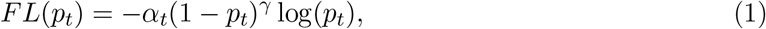

where *p*_*t*_ is the predicted probability of the true class, *α*_*t*_ is the class weight, and *γ* is the focusing parameter. In this study, class weights were set inversely proportional to class frequencies, and the focusing parameter was tuned, with an optimal value close to 3. This loss function helps the classifier concentrate on minority and hard examples.

### 3.5 Training Configuration

BioBERT was trained with the following configuration:

- **Optimizer:** AdamW
- **Learning rate:** 2 *×* 10^*−*5^
- **Batch size:** 8 (training and evaluation)
- **Number of epochs:** 4
- **Maximum sequence length:** 128 tokens
- **Early stopping:** based on validation F1-score

Early stopping on validation F1-score was used to avoid overfitting. Random seeds were fixed for all components (data splitting, model initialization, etc.) for reproducibility.

### 3.6 Evaluation Metrics

All models were assessed using accuracy, precision, recall, and F1-score.

## 4 Results and Discussion

This section presents the results of the proposed BioBERT-based model and compares its performance with the TF-IDF + SVM and TF-IDF + Random Forest baselines.

### 4.1 BioBERT Performance

The proposed BioBERT model achieves an overall accuracy of **85%**, a macro F1-score of **0.77**, and a weighted F1-score of **0.86**. Class-wise performance is summarized in Table 2.

**Table 2.** Class-wise performance of the BioBERT model.

| Class | Precision | Recall | F1-score |
| --- | --- | --- | --- |
| Safe (0) | 0.90 | 0.92 | 0.91 |
| Neutral (1) | 0.45 | 0.76 | 0.56 |
| Unsafe (2) | 0.93 | 0.78 | 0.85 |

The model performs consistently well on the majority classes while achieving a substantial improvement on the minority neutral class. The high neutral-class recall (0.76) demonstrates the effectiveness of the imbalance-aware training strategy. Fig. 1 shows the class-wise F1-scores across models.

**Figure 1.**
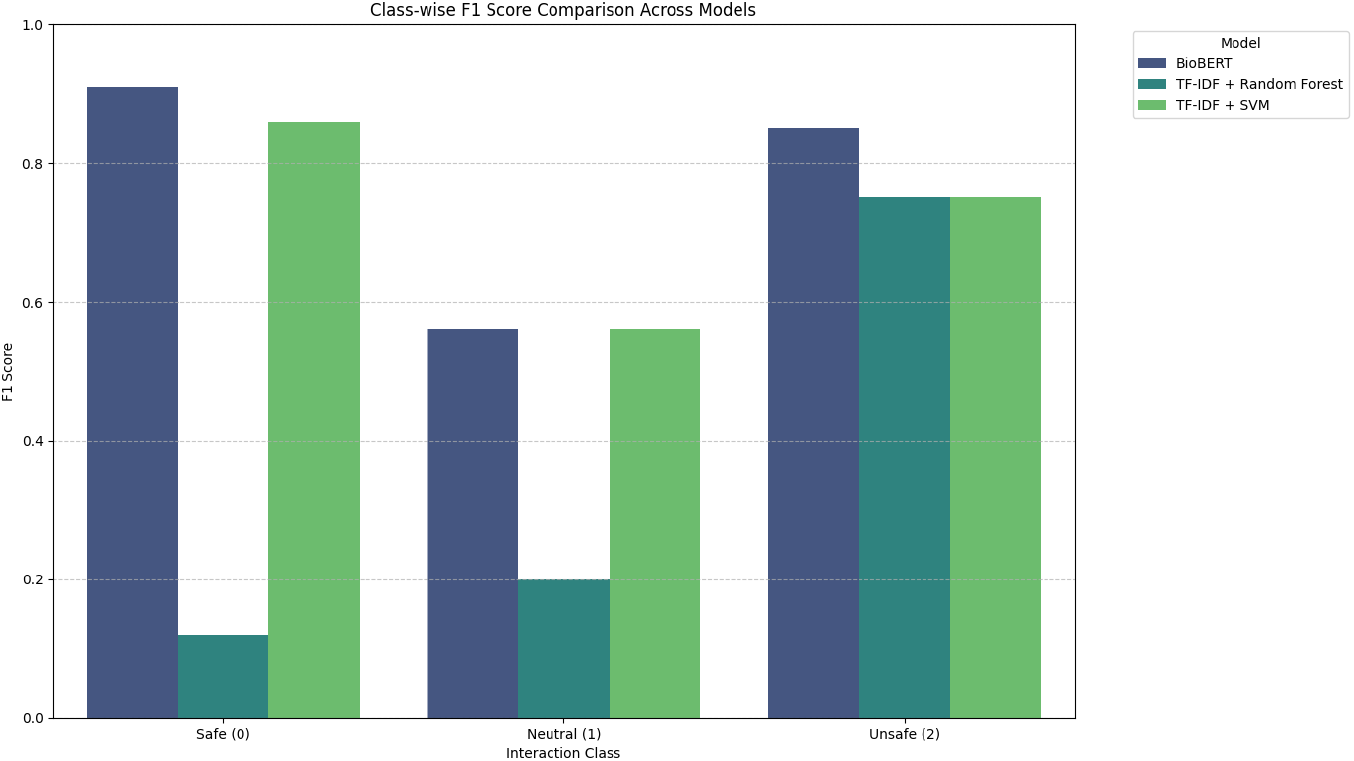
Class-wise F1-score comparison across models.

### 4.2 TF-IDF + Random Forest Performance

TF-IDF with Random Forest performs significantly worse than the proposed approach, with an overall accuracy of **39%** and a macro F1-score of **0.36**. The model is highly imbalanced in its behavior, with extremely low recall for the safe class and overfitting toward the minority class, indicating poor generalization.

### 4.3 TF-IDF + SVM Performance

TF-IDF with SVM performs better than Random Forest but still underperforms BioBERT, achieving an accuracy of **79%** and a macro F1-score of **0.72**. While it performs reasonably on majority classes, it does not fully capture contextual relationships in biomedical text.

### 4.4 Comparative Analysis

- BioBERT achieves the highest overall accuracy (85%) and macro F1-score (0.77).
- BioBERT substantially improves minority-class recall compared with the baselines.
- TF-IDF + SVM is a strong baseline but lacks contextual understanding.
- TF-IDF + Random Forest performs poorly owing to instability and sensitivity to class imbalance.

Fig. 2 further illustrates the comparison between BioBERT and the baselines.

**Figure 2.**
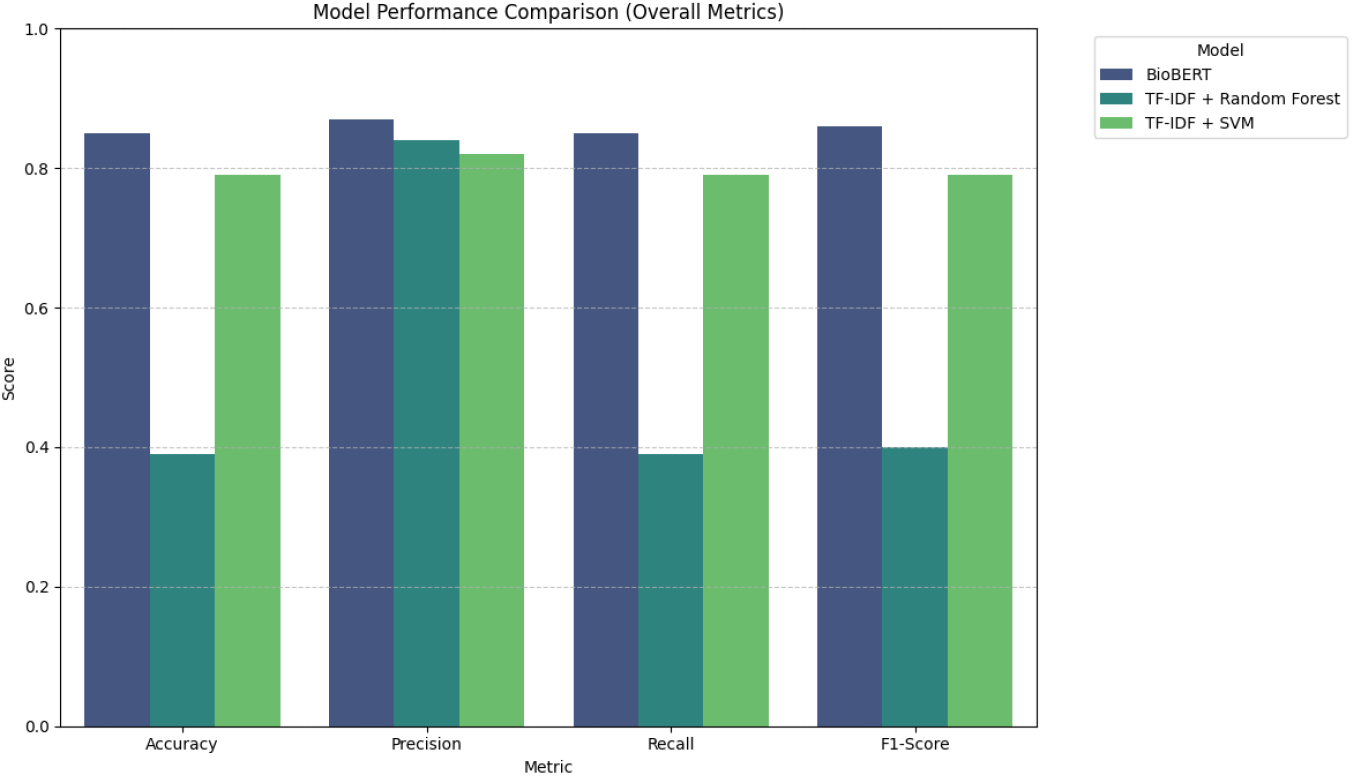
Model performance comparison across different metrics.

## 4.5 Discussion

The superior performance of BioBERT can be attributed to its domain-specific pretraining on biomedical corpora and its ability to model contextual dependencies. In addition, focal loss lets the model focus on difficult and underrepresented samples, improving minority-class performance.

**Figure 3.**
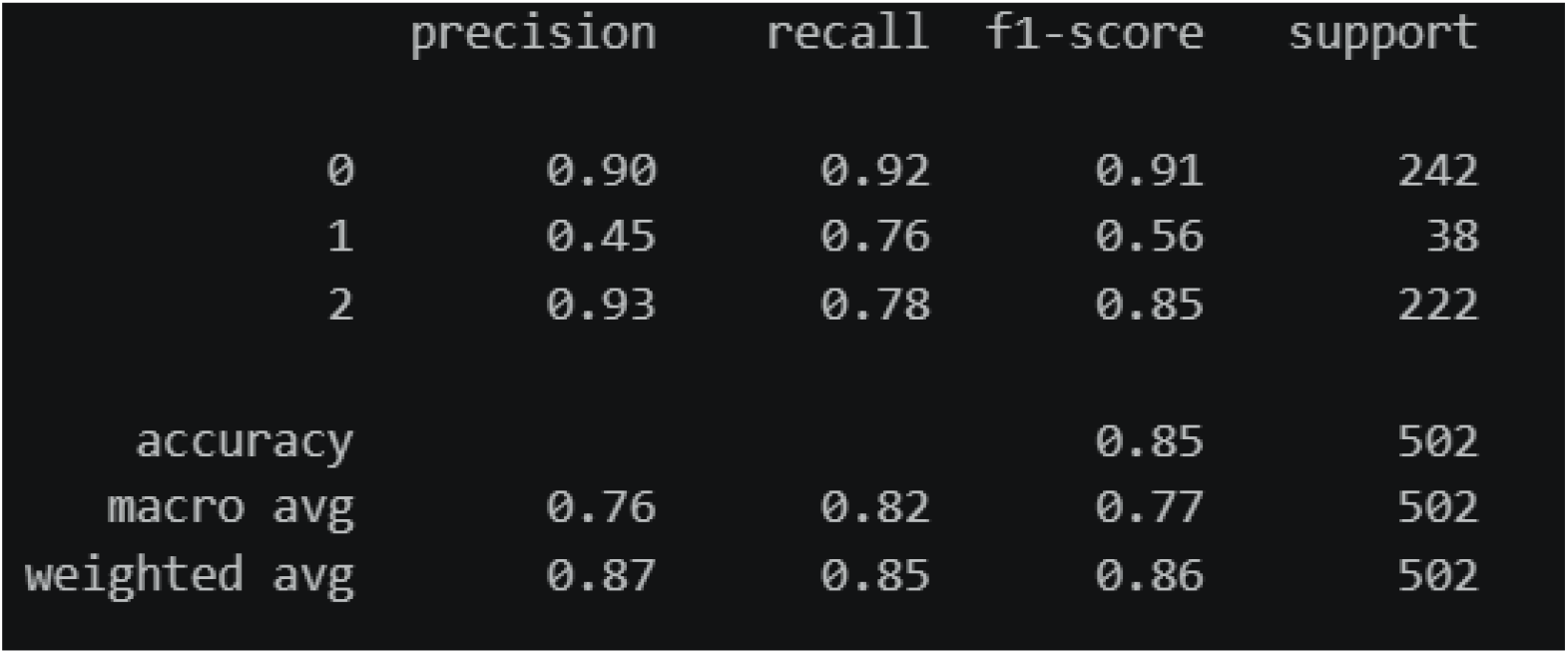
Classification report of the BioBERT model showing precision, recall, F1-score, and support for each class.

The confusion matrix in Fig. 4 illustrates the classification behavior of the model.

**Figure 4.**
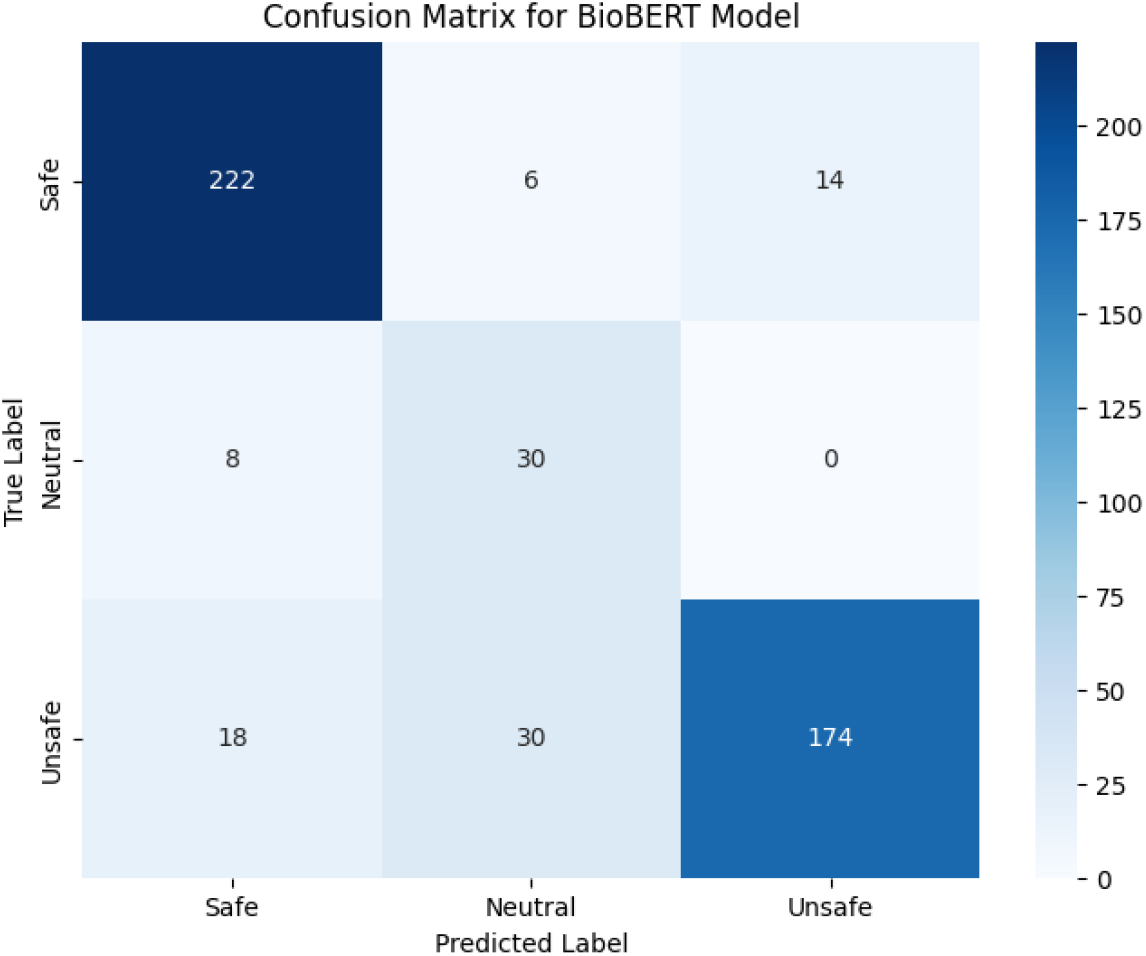
Confusion matrix of BioBERT model predictions.

However, the precision of the neutral class remains relatively low compared with the other classes, indicating that distinguishing neutral interactions remains challenging. This limitation is primarily due to the inherent ambiguity of such interactions and the limited number of samples.

Overall, the results suggest that combining transformer-based models with imbalance-aware optimization can enhance drug–food interaction classification.

## 5 Conclusion

We presented a framework for classifying DFIs based on contextual biomedical language models. The approach uses BioBERT to encode semantic relations in the biomedical domain and an imbalance-aware training method to address skewed class distributions. Experiments show high accuracy and F1-score, and a higher ability than conventional methods to handle minority classes.

The key contributions are a pipeline for converting unstructured drug–food interaction text into structured datasets, the use of BioBERT for DFI classification, the incorporation of focal loss, and a leakage-free evaluation protocol. Some challenges remain; in particular, the model predicts the neutral class less reliably, likely owing to the complexity of this class.

Future work includes using a larger biomedical text corpus, human-in-the-loop and semi-supervised labeling, newer deep learning architectures and ensembles, and external knowledge sources such as biomedical ontologies and knowledge graphs.

## Data and Code Availability

The dataset is publicly available on Kaggle [6], and the complete source code is available at https://github.com/NalgondaLokesh/Drug-Food-Interaction-BioBert.

## Acknowledgments

The authors thank the Department of Computer Science and Engineering at Neil Gogte Institute of Technology for their support.

## Author Contributions

